# Optical photothermal infrared spectroscopy (O-PTIR): a promising new tool for bench-top analytical palaeontology at the sub-micron scale

**DOI:** 10.1101/2024.02.08.579492

**Authors:** C.C. Loron, F. Borondics

**Affiliations:** UK Centre for Astrobiology, School of Physics and Astronomy, University of Edinburgh, Edinburgh, UK.; SMIS Beamline, Synchrotron SOLEIL, RD128 L’Orme des Merisiers, 91190, Saint-Aubin, France.

**Keywords:** Infrared Spectroscopy, O-PTIR, Biosignature, Rhynie Chert, Plant

## Abstract

The identification of preserved organic material within fossils is challenging. Well-established vibrational spectroscopy techniques, such as micro-FTIR (Fourier Transform Infra-Red spectroscopy), have been widely used to investigate organic fossils’ molecular composition. However, even when well-adapted to study objects several tens of micrometre across, they still suffer from limitations, notably regarding resolution and sample preparation requirements. Optical Photothermal Infrared Spectroscopy (O-PTIR), a recently developed technique, overcomes the challenges of bench-top FTIR spectroscopy. By combining an IR excitation laser with a 532 nm green probe laser, this technique allows molecular characterization at high spectral resolution (~2 cm^−1^) and with extremely fine spatial resolution (~500 nanometres). Additionally, problems linked with sample thickness, surface roughness and particle shape/size are mitigated when compared with FTIR or Atomic Force Microscopy-based nanoIR techniques. Here we show that O-PTIR can be used to easily and successfully map the molecular composition of small organic fossils preserved in silica matrix (chert) in petrographic thin sections. Our study reveals that O-PTIR resolves spatial heterogeneities in the preserved molecular composition of organic fossils (spores and plants) at a sub-micron scale, and that such heterogeneities occur in the cuticle in an early Devonian plant, where they suggest a structural organisation comparable to modern plants. These results on 400 million years old fossils, validate O-PTIR as a powerful and extremely promising new tool for nanoanalytical palaeontology.

---

Molecular signatures preserved in fossils constitute an important archive of the biology and ecology of ancient organisms (e.g., Briggs, 1995; De Leeuw et al., 2006). However, extracting relevant information is often complex due to the loss and transformation of compounds during decay and fossilization (Parry et al., 2018), the heterogeneities of preservation (Pang et al., 2021, Loron et al., 2022), the presence of contaminants (e.g., Rasmussen et al., 2021), and the limitations of analytical techniques.

Bench-top techniques like micro-Fourier Transform InfraRed (micro-FTIR) and Raman spectroscopy have provided great insights into the molecular composition and preservation of key cellular components in both micro- and macrofossils (e.g., Marshall et Marshall, 2015; Wiemann et al., 2020; McCoy et al., 2020; Loron et al., 2022; 2023) for resolution of several micrometres. Raman spectroscopy can achieve very high spatial resolution and provides complementary molecular information than FTIR but, with thermally mature organic material, is often limited to palaeothermometry (e.g., Baludikay et al., 2020). Benchtop FTIR analyses, on the other hand, remain limited by the spatial resolution of the apparatus. Nano Infrared (IR) techniques combining Atomic Force Microscopy (AFM) and infrared spectroscopy could offer an alternative to this spatial limitation by allowing studies of polymers at the scale of ~10 nm (Liu et al., 2021) but require complex sample preparation. High resolution approaches using synchrotron light can also overcome some of these limitations and reach higher spatial resolution (e.g., ~10 μm in Loron et al., 2022) to an extent defined by the diffraction of light and by the signal/noise ratio below diffraction limited resolution, restricting them to large material (e.g., fossils >100 μm).

Globally, sample preparation for spectroscopy is often challenging and may be hampered by the need to damage or destroy original material to conform to strict size, surface texture or thickness requirements or to bring a region of interest to the surface of a sample by removing overlying material. Many of these limitations are especially serious for work on precious palaeontological material such as museum specimens, for which destruction of sample (or even part of the sample), could be prohibited and not advisable to preserved geological heritage. Similarly, limitations due to spatial resolution, even for non-destructive approaches, can be problematic for the analyses of historical curated material depending on how these samples where prepared (use of organic resin, sample impregnation) because a large aperture involve more risk of acquiring signal from material surrounding the target. A high spatial resolution can facilitate the targeting of precise features and circumvent – or minimize – these acquisition artefacts.

Here we show the use a novel bench-top technique overcoming the limitations of the traditional spectroscopic approaches of fossil material: Optical-Photothermal infrared spectroscopy (O-PTIR). This technique allows very fine characterization in mid-infrared (3000-920 cm^−1^) of few 100 nm-size regions at the surface of a sample or a few microns below the surface (e.g., Freitas et al., 2021). The spectral resolution is also very high (2 cm^−1^). Thus, this technique overcomes most limitations of traditional FTIR spectroscopy in terms of sample preparation and spatial resolution. The O-PTIR infrared (IR) signal is obtained by the combination of a pulsed IR laser and a continuous wave visible laser (532 nm) (Figure 1). An IR spectrum is generated by measuring the photothermal excitation occurring when the IR laser matches an absorption mode in the sample (Zhang et al., 2016; Olson et al., 2020). Previously, O-PTIR was successfully used in the study of cultural heritage objects (e.g., Beltran et al., 2021; Marchetti et al., 2021) and biological samples (e.g., Klementieva et al., 2020; Ahn et al., 2022; Lima et al., 2022) but has not been applied to palaeontological material.

**Figure 1.**
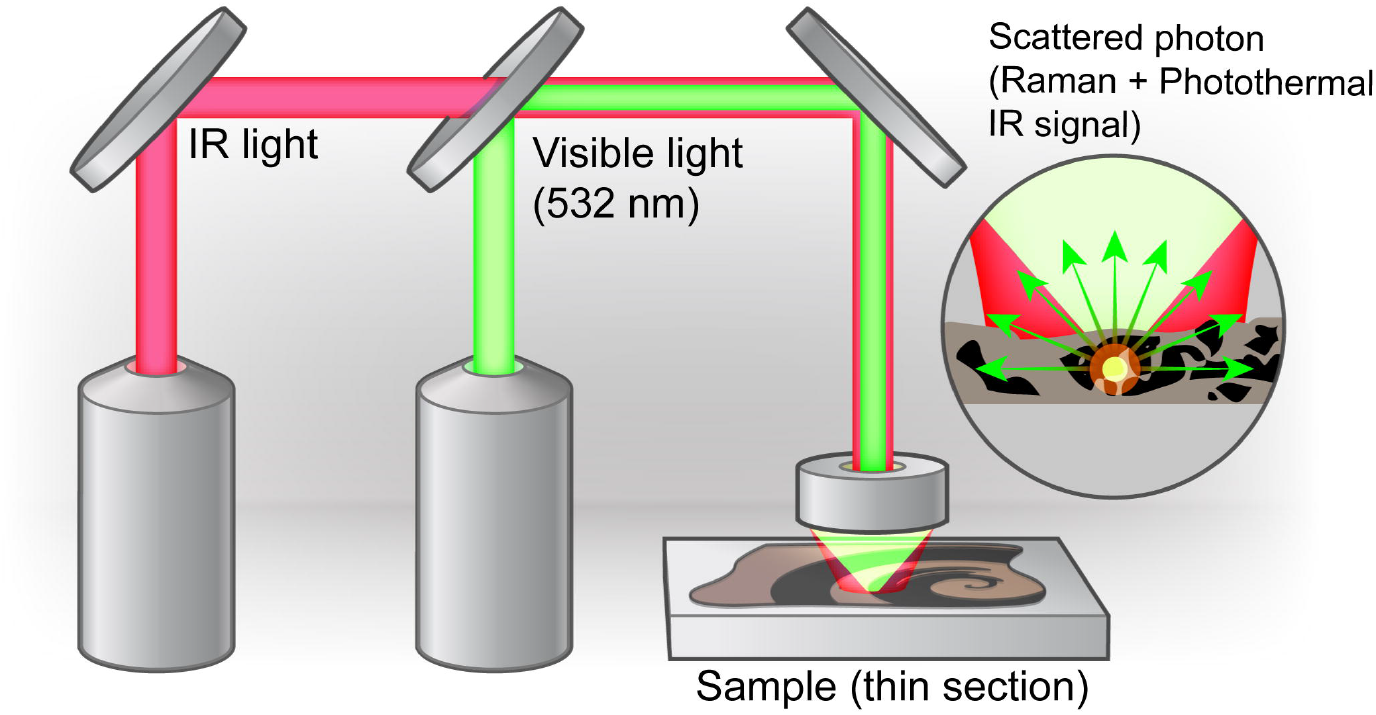
Schematic of optical photothermal infrared spectroscopy (O-PTIR). Partially adapted from Olson et al., 2020.

For this investigation, we applied O-PTIR on fossilized plant material from the exceptionally preserved 407 Ma Rhynie Chert assemblage from Aberdeenshire (Scotland) showing promise for the future studies of early land plants and their biotic milieu.

## Materials and methods

The studied fossils are from the 407 million years old Devonian Rhynie chert assemblage in Aberdeenshire, Scotland. This assemblage contains an important diversity of plant, fungi, animal, bacteria, and algae (Garwood et al., 2019) deposited in a terrestrial hot spring environment, and embedded in a silica matrix. The exceptional preservation state of these organisms makes the Rhynie chert an inestimable archive of early terrestrial biota during the *“greening of the land”* (Kerp et al., 2018).

We studied 2 fossils in a rock thin section (RF-f, Planetary Palaeobiology lab): one plant spore specimen and one specimen of the early land plant *Aglaophyton*, certainly the most common specimens of the Rhynie chert plants (Figures 2 and 3). *Aglaophyton* presents dichotomizing axis completed by spindle shape sporangia (the structures making and storing spores) and constitutes one of the earliest examples of fungal-plant mycorrhizal interactions (Edward, 1986; Remy and Hass, 1996; Kerp et al., 2018). The fossils are preserved in 3 dimensions in the chert matrix (silica).

**Figure 2.**
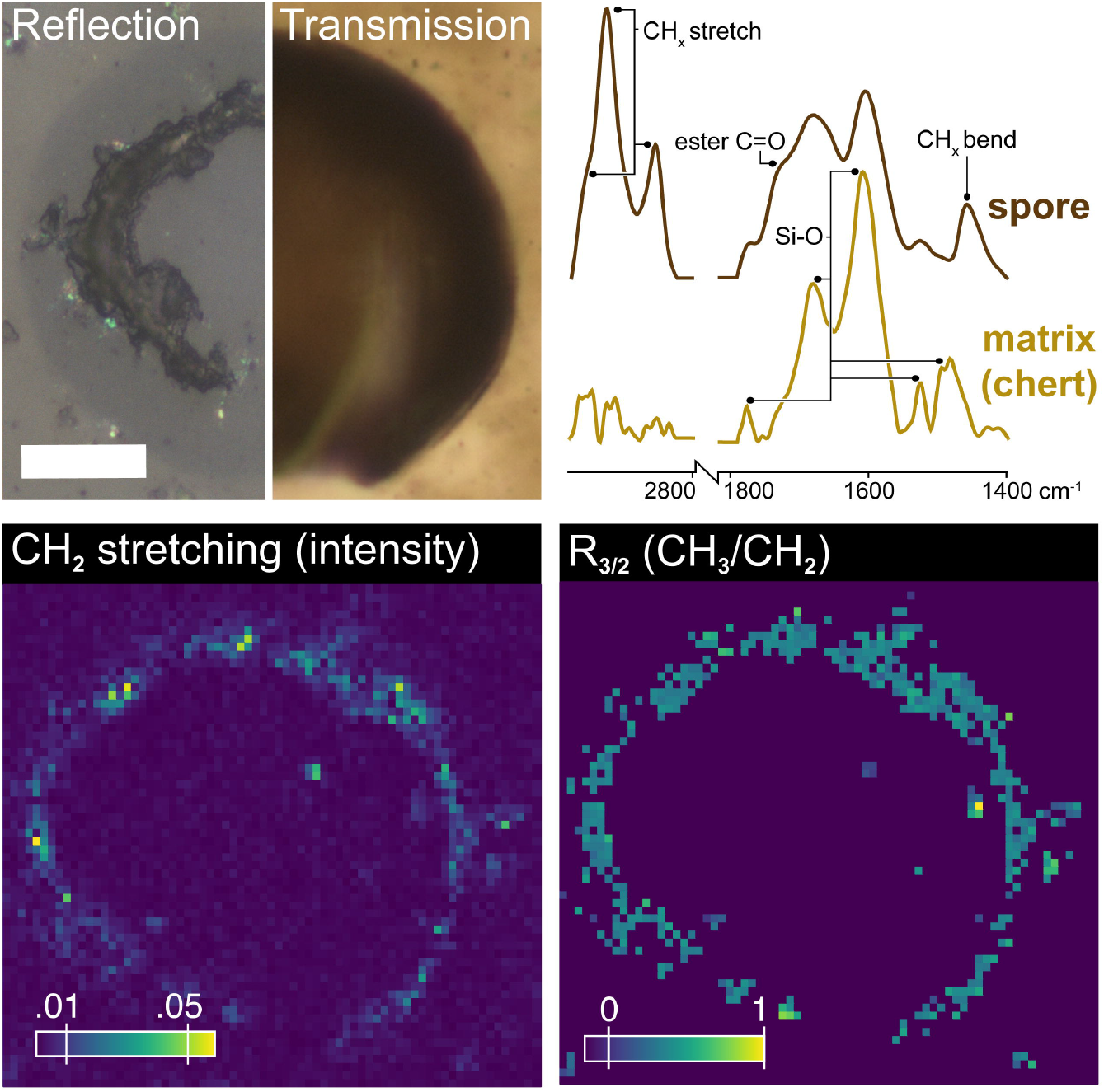
Spore specimen microphotographs in reflected and transmitted light (top left) and O-PTIR spectra of the spore tissue and adjacent matrix (top right). The intensity maps in the bottom illustrate the heterogeneities in intensity for the aliphatic groups (left) and for the R_3/2_ ratio map (right). Scale bar is 20 μm.

**Figure 3.**
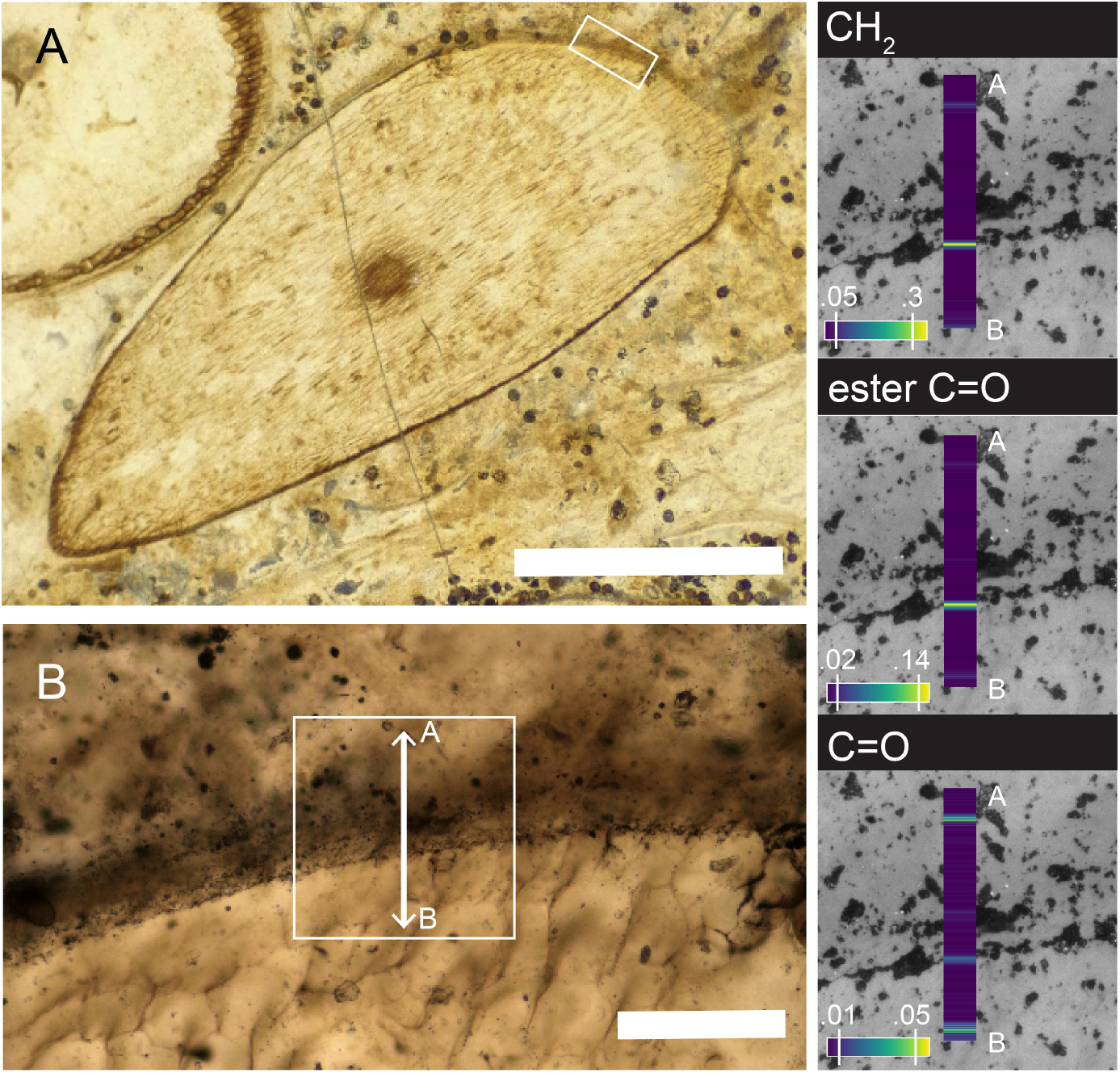
Studied *Aglaophyton* specimen (A) and zoom detail on the analysed cuticle zone (B). Intensity for several functional group (CH_2_, ester and C=O) for the transect A-B are shown on the right, superimposed on reflected light image (the width of the colour maps is exaggerated, the true width of the point is 0.5 μm). Scale bars is 2 mm in A and 200 μm for B.

The experiment was performed on a mIRage infrared microscope (Photothermal Spectroscopy Corp., Santa Barbara, CA) at the SMIS beamline (SOLEIL Synchrotron, Saint Aubin, France).

Spectra were acquired in reflection mode at a spectral resolution of 2 cm^−1^ with an IR laser power of 100%, probe laser power of 40% and detector gain of 50% using the PTIR Studio software. To avoid the overwhelming influence of Si-O and Si-C bond between 1400 and 900 cm^−1^, the spectra were acquired in the range 3000-1400 cm^−1^. Spectral pre-processing, including baseline correction and smoothing, were conducting in Quasar (Toplak et al., 2017; 2021).

The transect through the plant cuticle (Figure 3) represents 318 acquisitions with a 0.5 μm step size. Each spectrum is the average of 20 repetitions.

Intensity map of the spore specimen (70×61 μm) was acquired using a 1 μm step for a total of 4270 spectra. For each spectrum, we calculated the ratio CH_3_/CH_2_ (R_3/2_; Igisu et al., 2009) based on the intensity of the 2960 (asymmetric CH_3_ stretching) and 2925 cm^−1^ (asymmetric CH_2_ stretching) bands, respectively. This ratio displays lower values as the length of the chain increases and higher values as length decreases or branching increases (Lin and Ritz, 1993; Igisu et al., 2009).

## Results

### O-PTIR for high-resolution investigation of small palaeontological object

The spore specimen spectrum is characterized by a strong aliphatic absorption with characteristic bands of CH_x_ stretching between 3000 and 2800 cm^−1^, and CH_x_ deformation at ca. 1460 cm^−1^ (see Table 1). Absorption bands at 1705, 1630 and 1520 cm^−1^ are usually associated with carbonyl group and aromatic moieties in the spectra of sporopollenin (Yule et al., 2000). In our spectrum, the silica overtones, vibrating at similar wavenumber, are strong and these three absorptions bands are not observed, as showed on Figure 2 and Table 1.

**Table 1.** Band assignments. The letter ν corresponds to stretching vibration and δ to deformation. Numbers in column 1 correspond to bands in spectra in Figure 4.

|  | Spore | Plant | Band assignment | References |
| --- | --- | --- | --- | --- |
|  | cm<sup>-1</sup> |  |  |  |
| 1 | 2955 | 2956-2958 | Asymmetric $\nu$(CH<sub>3</sub>) | Coates, 2000 |
| 2 | 2928 | 2921-2925 | Asymmetric $\nu$(CH<sub>2</sub>) | Coates, 2000 |
| 3 | 2857 | 2853-2856 | Symmetric $\nu$(CH<sub>2</sub>) | Coates, 2000 |
| 4 | 1772 |  | Si-O | Ito and Nakashima, 2002 |
| 5 | 1727 | 1731-1733 | $\nu$(C=O) ester | Coates, 2002 |
| 6 | 1680 | 1687 | Si-O | Ito and Nakashima, 2002 |
| 7 | | 1668-16658 | $\nu$(C=O); amide I; $\nu$(C=C); $\nu$(C=N) | Coates, 2002; Movasaghi et al., 2008; Mohsin et al., 2018 |
| 8 | 1605 | 1609-1614 | Si-O | Ito and Nakashima, 2002 |
| 9 | | 1581 | $\delta$(N-H); COOH; carboxylate | Coates, 2002; Movasaghi et al., 2008; Lu and Miller, 2002 |
| 10 | | 1540 | $\nu$(C-N); $\delta$(N-H); COOH; carboxylate | Coates, 2002; Movasaghi et al., 2008; Lu and Miller, 2002 |
| 11 | 1525 | 1529 | Si-O | Ito and Nakashima, 2002 |
| 12 | 1496 |  | Si-O | Ito and Nakashima, 2002 |
| 13 | 1457 | 1465-1467 | $\delta$(CH<sub>2</sub>); $\delta$(C-H) | Coates, 2002; Movasaghi et al., 2008 |
| 14 |  | 1415-1418 | Si-O | Ito and Nakashima, 2002 |

The intensity maps reported for the spore specimen reveals light heterogeneities in intensity for the aliphatic groups (Figure 2). These variations could be due to differences in original aliphatic composition but their scattering around the specimen suggest they are more likely resulting from small local preservation differences or differences in signal intensity (e.g., due to small variation in sample topography). By converting the map to a R_3/2_ ratio map, we illustrate the actual change in aliphatic chain-length in the fossil wall, reflecting the change in fossil organic matter composition (Figure 2). The values for R_3/2_ within the spore wall vary between ca. 0.3 and 0.9. Such ratio values correspond to carbon chain containing between ca. 9 and 18 carbons (Igisu et al., 2009).

The transect acquired through the cuticle of the plant specimen (Figure 3) reveals variations of CH_x_ stretching, ester C=O and ester C=O absorptions, evidencing three different zones, for which each average spectra can be seen on Figure 4. Band assignments can be seen in Table 1. Interestingly, differences in the IR spectra resolve the cuticle of the plant into three layers. Spectra from all three show absorptions at ca. 1680, 1610, 1530, 1518 cm^−1^ corresponding to the overtone vibration of Si-O bonds (Ito and Nakashima, 2002). However, the spectra are dominated by the contribution of organic functional groups. The outer layer, in the exterior of the plant, is dominated by strong aromatic C=C absorption and a moderate aliphatic CH absorption. The middle layer shows an intense aliphatic absorption and strong ester C=O contribution. Finally, the inner layer, in the epidermis, is dominated by a high aliphatic absorption, contribution of ester C=O, C=O, carboxyl, carboxylate and N-moieties, attributed to condensation products of proteins and sugars (Wiemann et al., 2020; McCoy et al., 2021, Loron et al., 2023). Calculation of the R_3/2_ ratio for each of the spectra is 0.39 for zone A (outer), 0.11 for zone B (middle) and 0.35 for zone C (inner). These values can be converted into an equivalent chain-length (ignoring branching) of ca. 15, 30 and 18 carbons, respectively (Igisu et al., 2009).

**Figure 4.**
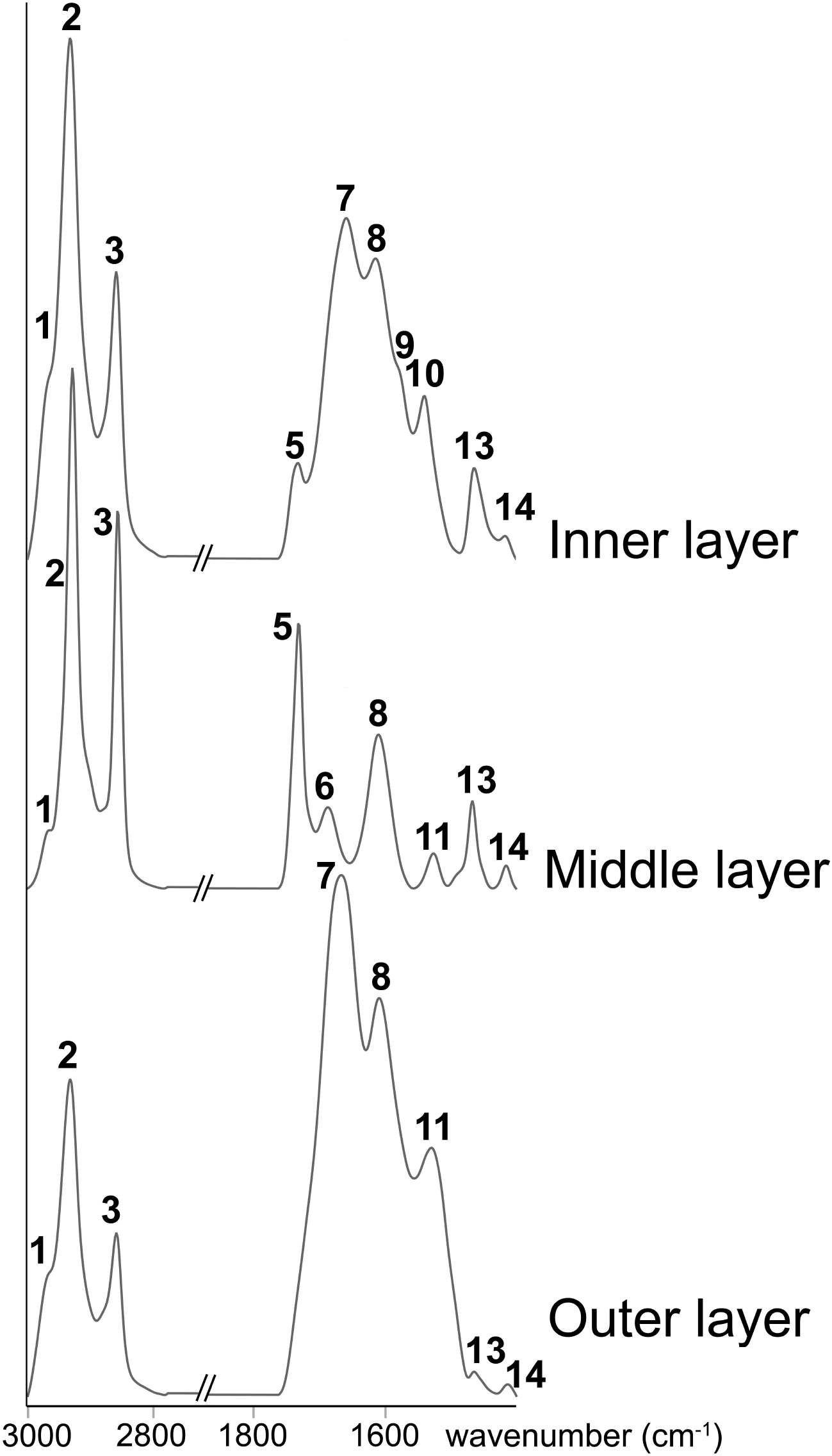
O-PTIR spectra for the three detected zone of the transect shown on Figure 3. Numbers correspond to band numbers in Table 1.

## Discussion and conclusion

Previous spectroscopic investigations of Rhynie Chert material provided important insights into the molecular preservation of its iconic fossils (Preston and Genge, 2009; Qu et al., 2015) and most recently have revealed that taxon-specific chemical fingerprints can be identified by ATR-FTIR spectroscopy (aperture size 100 μm; Loron et al., 2023). Here, we show that O-PTIR can resolve molecular differences within organisms and tissues at a much smaller spatial scale. As shown by Figure 2, the infrared signature of the organic content of a fossil plant spore ca 80 μm in diameter can be mapped at very high level of resolution (step of 1 μm), revealing small heterogeneities in aliphatic absorption and chain length (R_3/2_ ratio map). This level of precision for such a small fossil is superior to what is usually accomplished even with synchrotron-FTIR (Igisu 2017, 2022; Loron et al., 2022).

The Rhynie Chert assemblage records several well studied plant taxa (e.g., *Asteroxylon, Aglaophyton, Rhynia, Horneophyton* and *Nothia)*. In all these taxa, the plant epidermis is covered by a well-developed cuticle (Kerp et al., 2018) but the composition and structure of these cuticles remain unclear. The prevailing model considers cuticle as a cutin matrix with embedded intracellular waxes and phenolic compounds, an innermost polysaccharide region and an outermost layer of epicuticular waxes, although the structure and composition vary between plant species (Fernández et al., 2016). Fossils might be expected to show structural organization similar to modern plant cuticle given the preservation potential of its different components, but fossils older than the Cenozoic (before 66 Ma) have hitherto failed to record this structure and rather show a dominance of aliphatic signal (Gupta et al., 2006). The present results obtained on *Aglaophyton* shows that the cuticle of this plant has been modified diagenetically into at least three layers with distinct molecular composition, supporting the morphological observations (Remy and Hass, 1996). The molecular characteristics of each of these zones are consistent with cuticular waxes, cutin and polysaccharide-cutin mix (Herredia-Guerrero et al., 2014), respectively. Although it cannot be excluded that post-depositional condensation of external material might have contributed to the observed molecular signal (e.g., aliphatic material, see De Leeuw et al., 2006; Gupta et al., 2006), the similarity between composition of these three distinct layers and their extant counterparts is striking and suggests they derived from waxes, cutin and polysaccharide precursors, modified by taphonomy, diagenesis and fossilization. Our results show that O-PTIR can successfully detect molecular ultrastructure within plant fossils, and that these fossils possessed a cuticular organisation and composition similar to modern plant taxa.

The ultra-high resolution and simplicity of sample preparation offered by O-PTIR allow the technique to insert well within palaeontological workflows, especially as complement to traditional infrared spectroscopy techniques, such as transmission or ATR-FTIR (Figure 5), but also as an alternative to these approaches when they are not applicable. Indeed, beyond visual inspection of samples with optical microscopy, the array of analytical approaches available are often dependant of the sample preparation technique (Wacey et al., 2017). For example, organic fossil and material extracted from the rock matrix using acid maceration (HF/HCl) can be deposited on silicon nitride (Si_3_N_4_) membrane windows for X-ray spectroscopy (Sforna et al., 2022). Si-N functional groups from the membrane window, around 800-1000 cm^−1^ (Lattemann et al., 2003), will slightly interfere with the organic signal from the fossils in transmission FTIR, an otherwise very well-adapted type of acquisition for the study of extracted organic microfossils (e.g., Steemans et al., 2010; Loron et al., 2022). By using reflection geometry, O-PTIR can circumvent this limitation allowing the samples to be preliminarily studied with IR before destructive X-ray fluorescence or XANES. Similarly, because it requires very few amounts of material and can be operated on various sample thickness, O-PTIR is well adapted to analyse historical or museum-based material for which authorized destructive sample preparation is often prohibited or drastically limited to thin-sections or small chips (Figure 5). Although adapted for double-polished and standard uncovered sections, O-PTIR analyses of fossils in covered thin sections would be drastically hindered by the glass cover, as silica would interfere with wavenumber below 1550 cm^−1^ (Ito and Nakashima, 2002).

**Figure 5.**
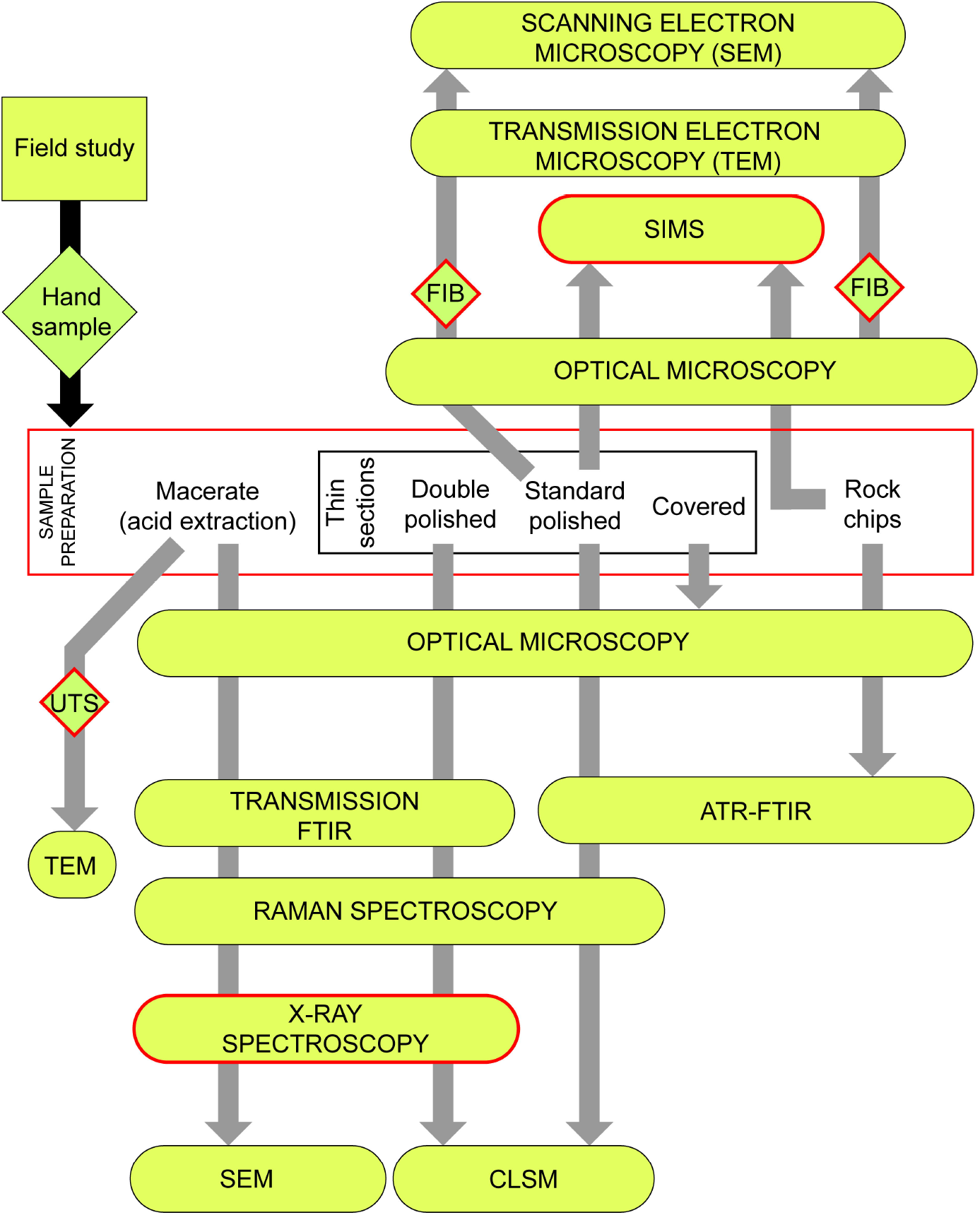
Examples of palaeontological workflows for the study of organic fossils. Red borders indicate destructive steps and measurements. O-PTIR can insert after any visual inspections and preferably before destructive measurements. For details and examples for each technique see Wacey et al. (2017). FIB, Focus Ion Beam section. UTS, Ultra-thin Section. ATR-FTIR, Attenuated Total Reflectance-FTIR. SIMS, Secondary Ion Mass Spectrometry. CLSM, Confocal Laser Scanning Microscopy.

Characterization of morphological architectures, elemental composition and crystallinity of fossil walls can be achieved using TEM on ultra-thin section on extracted fossils (Ohler, 1977; Moczydlowska and Willman, 2009; Taylor and Strother, 2009; Guignard, 2019) or by directly obtaining sections from the host rock using a focus ion beam (FIB), which allow targeting of individual microfossil or specific parts of individual microfossils (e.g., Schiffbauer and Xiao, 2011). The high spatial resolution of O-PTIR has the potential to target area of hundreds of nm within ultra-thin and FIB sections. Complementing TEM morphological and elemental characterisation, this offers a promising perspective for the study of architectural compositional differences.

In summary, O-PTIR can easily be inserted in palaeontological workflows, practically before a destructive measurement, in addition to be compatible with a wide range of sample preparations (extracted microfossils, rocks chips, uncovered thin-sections, FIB and ultrathin sections). Altogether, the presented results introduce O-PTIR as a novel, quick, non-destructive, extensively promising, and extensible approach for the molecular imaging and characterisation of palaeontological material.

## Acknowledgment

This research was supported by The Royal Society (UK), the Belgium Wallonia Brussels programme WBI.WORLD and the Leverhulme Trust (C.C.L.). We thank S. McMahon at University of Edinburgh for the access to the sample and useful discussions during the writing of this manuscript. I. Febbrari (University of Edinburgh) is thanked for the preparation of the thin section. SOLEIL Synchrotron (Saint Aubin, France) is acknowledged for providing access to the mIRage instrument at the SMIS beamline (proposal 99220020).

